# Ventral hippocampus mediates inter-trial responding in signaled active avoidance

**DOI:** 10.1101/2024.03.18.585627

**Authors:** Cecily R. Oleksiak, Samantha L. Plas, Denise Carriaga, Krithika Vasudevan, Stephen Maren, Justin M. Moscarello

**Affiliations:** Department of Psychological and Brain Sciences, Texas A&M University, College Station, TX 77845; Texas A&M Institute for Neuroscience, Texas A&M University, College Station, TX 77845; Department of Psychological Science, University of Texas Rio Grande Valley, TX 78539

**Keywords:** Ventral hippocampus, Inter-trial Intervals, Avoidance, Context, DREADDs

## Abstract

The hippocampus has a central role in regulating contextual processes in memory. We have shown that pharmacological inactivation of ventral hippocampus (VH) attenuates the context-dependence of signaled active avoidance (SAA) in rats. Here, we explore whether the VH mediates intertrial responses (ITRs), which are putative unreinforced avoidance responses that occur between trials. First, we examined whether VH inactivation would affect ITRs. Male rats underwent SAA training and subsequently received intra-VH infusions of saline or muscimol before retrieval tests in the training context. Rats that received muscimol performed significantly fewer ITRs, but equivalent avoidance responses, compared to controls. Next, we asked whether chemogenetic VH activation would increase ITR vigor. In male and female rats expressing excitatory (hM3Dq) DREADDs, systemic CNO administration produced a robust ITR increase that was not due to nonspecific locomotor effects. Then, we examined whether chemogenetic VH activation potentiated ITRs in an alternate (non-training) test context and found it did. Finally, to determine if context-US associations mediate ITRs, we exposed rats to the training context for three days after SAA training to extinguish the context. Rats submitted to context extinction did not show a reliable decrease in ITRs during a retrieval test, suggesting that context-US associations are not responsible for ITRs. Collectively, these results reveal an important role for the VH in context-dependent ITRs during SAA. Further work is required to explore the neural circuits and associative basis for these responses, which may be underlie pathological avoidance that occurs in humans after threat has passed.

## 1. Introduction

Proactive avoidance strategies can be highly adaptive to the extent that avoidant behavior prevents a harmful outcome [1,2]. However, an avoidance response that continues after danger has passed does not provide the same adaptive benefit and can even become maladaptive if it interferes with the resumption of preferred patterns of behavior [3–5]. Indeed, persistent forms of avoidance that disrupt normal life activities are a unifying symptom of disorders related to fear, anxiety, and obsessive thinking. Relatively little is known about the neural and behavioral processes underlying avoidant behaviors that persist despite being uncoupled from their protective/preventative effects. To explore this phenomenon, we used a signaled active avoidance (SAA) procedure in which persistent inter-trial responses (ITRs) are elicited by discrete conditioned stimuli (CSs).

Previous studies suggest that ITRs in SAA are driven by aversive associations with the environment in which training occurs [6]. Indeed, a similar role for context in the production of ITRs has been demonstrated for appetitive instrumental behavior [7]. The ventral hippocampus (VH) and associated cortices are important substrates for aversive contextual processes, such as the acquisition and expression of aversive contextual associations [8–11], the contextually-mediated renewal of previously-extinguished aversive CSs [12–15], and context-dependent instrumental avoidance [16–18]. VH is also required for classic anxiety-like behaviors, in which animals organize their exploratory behavior to minimize contact with threatening aspects of an experimental apparatus [19–23]. These results are consistent with the proposal that VH contains a map that links interior representations of aversive states with environmental stimuli [24]. We hypothesized that an aversive association with the SAA training context engages VH to produce ITRs.

Consistent with this hypothesis, our results demonstrate that pharmacological inactivation of VH suppresses ITRs while having no effect on avoidance responses (ARs) in the training context, and that chemogenetic excitation of VH neurons facilitates the expression of ITRs but not ARs in both the training and alternate contexts. Though these data reveal the contribution of VH to ITR expression, we also show that extinction of the SAA training context had no significant effect on ITRs, suggesting that aversive associations with the training environment do not mediate ITR expression. This result indicates that an alternative associative process, such as a residual CS-elicited aversive state that lingers beyond performance of the avoidance response [25], engages VH to drive the expression of ITRs in SAA.

## 2. Methods

### 2.1 Subjects

Subjects were 221 experimentally naïve, adult Sprague-Dawley rats (123 males, 98 females) weighing 200-300 g at time of arrival. Of these, 25 males contributed data to a previously published study (Oleksiak et al., 2021). The remaining 98 male and 98 female subjects comprised equally divided, mixed-sex cohorts for all other experiments. Rats were purchased from Envigo for Experiment 1, from Envigo and Charles River for Experiment 2, from Charles River for Experiment 3, and Envigo for Experiment 4. Multiple vendors were used to accommodate availability issues stemming from COVID-related changes in demand for experimental subjects. Rats were individually-housed in the Interdisciplinary Life Sciences Building vivarium and maintained on a 14h light: 10h dark cycle beginning at 7 AM. All subjects had *ad libitum* access to food and water. All experiments were conducted during the light cycle. Experimenters gently handled each rat for two minutes/day for four days prior to each experiment. All experiments were approved by the Animal Care and Use Committee from Texas A&M University.

### 2.2 Surgery

For experiments involving muscimol inactivation of ventral hippocampus (VH), surgical methods have been published previously (Oleksiak et al., 2021). This approach was adapted for experiments involving DREADD activation of VH neurons. Briefly, rats were anesthetized with isoflurane (5% for induction, slowly reduced throughout the procedure) and fixed in a stereotaxic apparatus. Lidocaine was injected subcutaneously at the incision site, and rimadyl was injected at the beginning of surgery for post-surgical analgesia. Then an incision was made in the scalp, and bore holes were placed to allow intracranial injections. Rats then received bilateral infusions of either AAV8-CAMKIIa-hM3D(Gq)-mCherry or AAV8-CaMKIIa-GFP (5×10^12^ vg/mL for both viruses; Addgene) into the VH (either A/P: -5.25 or -5.65, M/L: ± 5, D/V: either -7.3 or -8.5 from bregma). Infusions were made at a rate of 0.1 µl/min for a total volume of 0.75 µl. Five minutes were allowed for diffusion after each injection. Injectors were then slowly removed at a rate of 0.1 mm/min for the next five minutes. Following surgery, subjects were allowed to recover for at least four weeks before tests.

### 2.3 Behavioral Apparatus

Six identical shuttle boxes (50.8 x 25.4 x 30.5 cm, LxWxH; Coulbourn Instruments) constructed of Plexiglas and metal were used for all training and testing. Each shuttle box was divided into two equal compartments (8 x 9 cm, WxH) between which subjects could pass freely through an aperture in the divider. The floors were made of conductive stainless-steel bars through which electric shock was delivered. Two speakers, one on either side of the length of each chamber, delivered a 2 kHz, 80 dB tone CS for a maximum of 15 seconds. A scrambled shocker delivered a 0.7 mA, 0.5 second footshock US through the floor. Each chamber was contained inside a larger sound-attenuating chamber. The subjects’ motion was monitored by two infrared arrays on either side of the divider between compartments. CS and US presentation were controlled by GraphicState Software (Coulbourn Instruments), which also collected shuttling data via the infrared arrays in each compartment

Two distinct shuttle-box contexts were created using methods published previously (Oleksiak et al., 2021). In all experiments, SAA training was conducted in the shuttle boxes with the house light off, black construction paper wall inserts with glow-in-the-dark stars, a 1% ammonia odor, and open doors of the sound-attenuating chamber during training (room lights also remained off). Rats were transported in black transport boxes without bedding. To create an alternate shuttle-box context, the house light in each compartment was illuminated (0.5 W light bulb), black-and-white striped wall inserts were placed behind the Plexiglas walls of the chamber, the box was scented with a 3% acetic acid solution, a black Plexiglas floor was placed over the conductive bars (no shocks were delivered in the alternate context), and the doors of the sound-attenuating chamber remained closed. Rats were transported to this alternate context in white transport boxes with bedding.

For locomotor activity testing (Experiment 2, see below) and two extinction-learning control conditions (Experiment 4, see below), five identical behavioral chambers (30 x 24 x 21 cm, L x W x H; Med-Associates, St Albans, VT) were used. Each box was a single compartment made of Plexiglas and metal. The floors were comprised of stainless-steel bars covered with a black Plexiglas floor insert. To fully distinguish this environment from the shuttle box training context, the alternate context cues were used in these single-chamber boxes (high-contrast striped backgrounds were placed behind the Plexiglas, the chamber was scented with acetic acid, and subjects were transported from the colony to the single-chamber boxes via white plastic carriers with bedding). A grating in one of the aluminum sidewalls allowed the delivery of tone CSs (2 kHz, 80 dB tone) from a speaker mounted on the outside of the box. Each box sat on a load-cell platform that recorded how the chamber was displaced by the rat’s locomotion. The data was collected by Threshold Activity software (Med-Associates). Each chamber’s load-cell voltages were digitized at 5 Hz, which provided one observation every 200 ms for locomotion measures.

### 2.4 Signaled Active Avoidance (SAA) Training & Test

SAA training and tests occurred in identical Coulbourn shuttle boxes. To ensure that all subjects associated tone with shock, the first trial of the first session of SAA training was comprised of a CS paired with a US that could not be avoided by shuttling. The rest of training consisted of avoidance trials, in which shuttling (crossing through the divider to the opposite compartment of the shuttle box) during the CS resulted in the immediate offset of the CS and the omission of the US. Shuttling during the CS was defined as an avoidance response (AR). Each SAA training session involved 30 such avoidance trials. Between each trial was an inter-trial interval (ITI) that averaged 120 seconds. Shuttling during the ITI was defined as an inter-trial response (ITR). Rats were considered poor avoiders and excluded from further analysis if they avoided on ≤ 20% of trials averaged for the last 3 days of training.

Tests of SAA were conducted under extinction conditions in either the training or alternate shuttle box context described above. During these tests, ten 15-sec CSs were presented separated by a 2-min ITI. CSs did not terminate with shuttling and no USs were presented.

### 2.5 Drug Infusions

Procedures for muscimol (GABA_A_ agonist) infusions into VH have been described previously (Oleksiak et al., 2021). Clozapine-N-oxide (CNO) was purchased from RTI International. For DREADD experiments, CNO was dissolved in 2.5% DMSO until clear liquid was achieved and then diluted in sterile saline to a concentration of 3 mg/kg/mL immediately before injection. Approximately 30 minutes before start of test or locomotion session, rats received i.p. CNO or 2.5% DMSO saline (vehicle) injections in the vivarium and were placed back in their home cage until transport to the shuttle boxes or locomotor test boxes.

### 2.6 Experimental procedures

#### 2.6.1 Experiment 1: Effect of pharmacological inactivation of VH on ITRs in the training context

Rats received chronic guide cannula implanted in VH and were allowed to recover before undergoing four days of SAA training. Following training but before testing, all subjects were assigned to treatment-order conditions. A pair of counterbalanced SAA tests were then conducted, one in the training and one in the alternate shuttle-box context. Both were conducted under extinction conditions (see above). Prior to each test, all subjects received intra-VH infusions of muscimol or vehicle. Here, we report novel ITR data in the training context from this experiment – AR data have been reported previously (Oleksiak et al., 2021).

#### 2.6.2 Experiment 2. Effect of chemogenetic activation of VH principal neurons on ITRs in the training context

Rats were injected with a virus bearing the gene construct for either the excitatory hM3Dq DREADD or GFP and were allowed to recover for at least three weeks. After recovery, rats underwent four days of SAA training. Treatment-order conditions for SAA testing were determined by balancing groups based on avoidance performance during training. Rats then underwent two days of SAA testing under extinction conditions. Both tests occurred in the training context. Thirty minutes prior to each, both hM3Dq and GFP groups received i.p. CNO or vehicle in a counterbalanced order. One week after testing, a subset of animals from both hM3Dq and GFP groups (7 hM3Dq and 6 blank-GFP rats) underwent an off-baseline test of locomotor activity in a novel environment (cubicle Plexiglas-and-metal chambers situated on load cells for the quantification of locomotion). These rats received two counterbalanced locomotor tests preceded by CNO or Vehicle, in which where they were allowed to move freely in the novel environment for 10 minutes.

#### 2.6.3 Experiment 3. Effect of chemogenetic activation of VH principal neurons on ITRs in an alternate shuttle-box context

All rats received VH viral infusions for the expression of hM3Dq or GFP at least three weeks before the start of SAA training. Most of the animals from both hM3Dq and GFP groups (32 hM3Dq and 27 GFP) received a cannula implantation procedure following the VH viral injection but prior to SAA. After four days of SAA training, these subjects received intracranial infusions and a pair of SAA tests (data not presented). Cannulated subjects then received an additional day of SAA training to re-establish baseline avoidance, followed by two SAA tests in the alternate shuttle-box context preceded by i.p. CNO or Vehicle in a counterbalanced order. For a subset of animals (7 hM3Dq and 8 GFP), testing in the alternate SAA context began following the four days of SAA training. All other elements of SAA testing were the same in both subsets of rats. Because ANOVA revealed that the additional procedure and training did not influence the systemic effects of CNO on ITR expression in hM3Dq and GFP subjects, we did not consider this variable in subsequent analyses.

#### 2.6.4 Experiment 4. Effect of context extinction on ITRs

Rats received 4 days of SAA training, as above. This was followed by three additional days under one of the following four conditions: 1) context extinction via exposure to the SAA training context in the absence of the CS or US, for the normal duration of a training session (73 minutes); 2) equivalent-duration exposure to single-chamber alternate context, also in the absence of the CS or US; 3) equivalent-duration exposure to single-chamber alternate context, with the delivery of 30 CSs/day separated by a 2-min ITI; and 4) equivalent-duration exposure to white 5-gallon buckets (handling control). Following three days of exposure to these conditions, all rats received a SAA test under extinction conditions, conducted in the SAA training context (as described above).

### 2.7 Histology

Rats in pharmacological and chemogenetic experiments were injected with a fatal dose of sodium pentobarbital (Fatal Plus; 100 mg/kg, Vortech Pharmaceuticals) and were transcardially perfused first with refrigerated saline followed by 10% formalin. The brains were collected and were left in 10% formalin for 24 hours. The brains were then transferred to 30% sucrose in PBS at 4°C until they sank. Brains were sliced coronally on a cryostat (-20°C) at 30 micron thickness. Slices were mounted on subbed slides and coverslipped with fluoromount to visualize the mCherry or GFP from the viral manipulation.

### 2.8 Data Analysis

ANOVAs followed by the appropriate *post hoc* tests were performed on all data. All statistical analyses were run in Statview version 5.0.1 (SAS Institute) in a MacOS 9 open-source emulator.

For all experiments, a poor-avoider criterion was applied following SAA training but prior to test. Animals averaging six or fewer ARs during the last three sessions of training were determined to be poor avoiders [26–28]. Because these experiments were designed to explore neural substrates in animals that successfully express SAA responses, poor avoiders were removed from our analyses.

## 3 Results

### 3.1 Experiment 1: Pharmacological inactivation of VH reduces the expression of ITRs

We first assessed whether VH is necessary for the expression of ITRs in SAA. Rats acquired ARs over the course of SAA training, prior to being assigned to drug (n=13) or vehicle (n=12) groups. Then, all subjects received SAA tests under extinction conditions in the training and alternate SAA contexts, preceded by intra-VH infusions of muscimol or saline, depending on group (behavioral procedure schematized in FIG 1A, only data collected from the SAA test in the training context are depicted here; all other AR data shown in: Oleksiak et al., 2021). As previously reported, VH inactivation had no effect on ARs (capped 1/CS) during tests conducted in the SAA training context (Oleksiak et al., 2021; FIG 1B, left). We also performed a novel analysis of the effects of VH inactivation on ITRs during the same test. A two-way ANOVA of mixed design with a within-subjects factor of ITI and a between-subjects factor of Drug (Muscimol or Vehicle) revealed a significant ITI X Drug interaction [*F(*9,207) = 2.232, *p* = 0.0213]. Thus, muscimol inactivation of VH suppresses ITRs while having no effect on AR expression. We speculate that vehicle and muscimol ITRs converge later in the session due to within-session extinction effects.

**Fig. 1.**
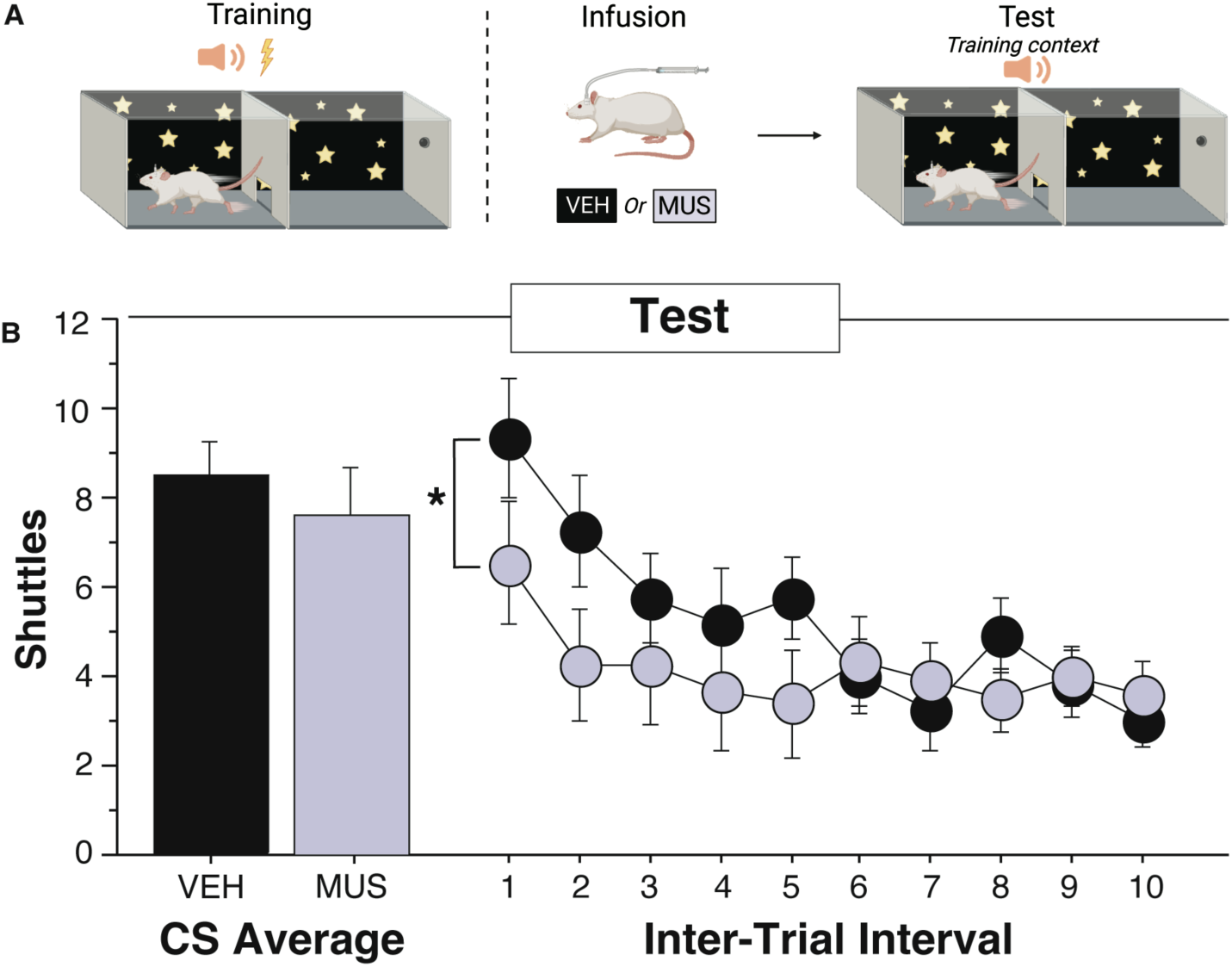
VH inactivation decreases ITI shuttling over the course of a 2-way SAA test in the training context. A) Rats were trained in two-way signaled active avoidance and had either muscimol (n=13) or vehicle (n=12) infused into the VH immediately before a test in the training context. Created with BioRender.com. B) The effect of muscimol and vehicle on capped CS shuttles and total ITI shuttles throughout the training context test. Muscimol significantly decreased the number of ITI shuttles during this test (* indicates p<0.05). All data calculated as means ± SEM. Adapted from Oleksiak et al., 2021.

### 3.2 Experiment 2: Chemogenetic activation of VH principal neurons drives expression of ITRs in the training context

Next, we sought to determine whether engagement of VH principal cells is sufficient to enhance ITR expression. To establish this, we infused a virus containing the gene construct for the excitatory hM3Dq DREADD or a blank-GFP control (AAV8-CamKII-hM3Dq-mCherry, n=14; or the control virus AAV8-CamKII-GFP, n=10). Representative images and viral spread are depicted in FIG 2A.

**Fig. 2.**
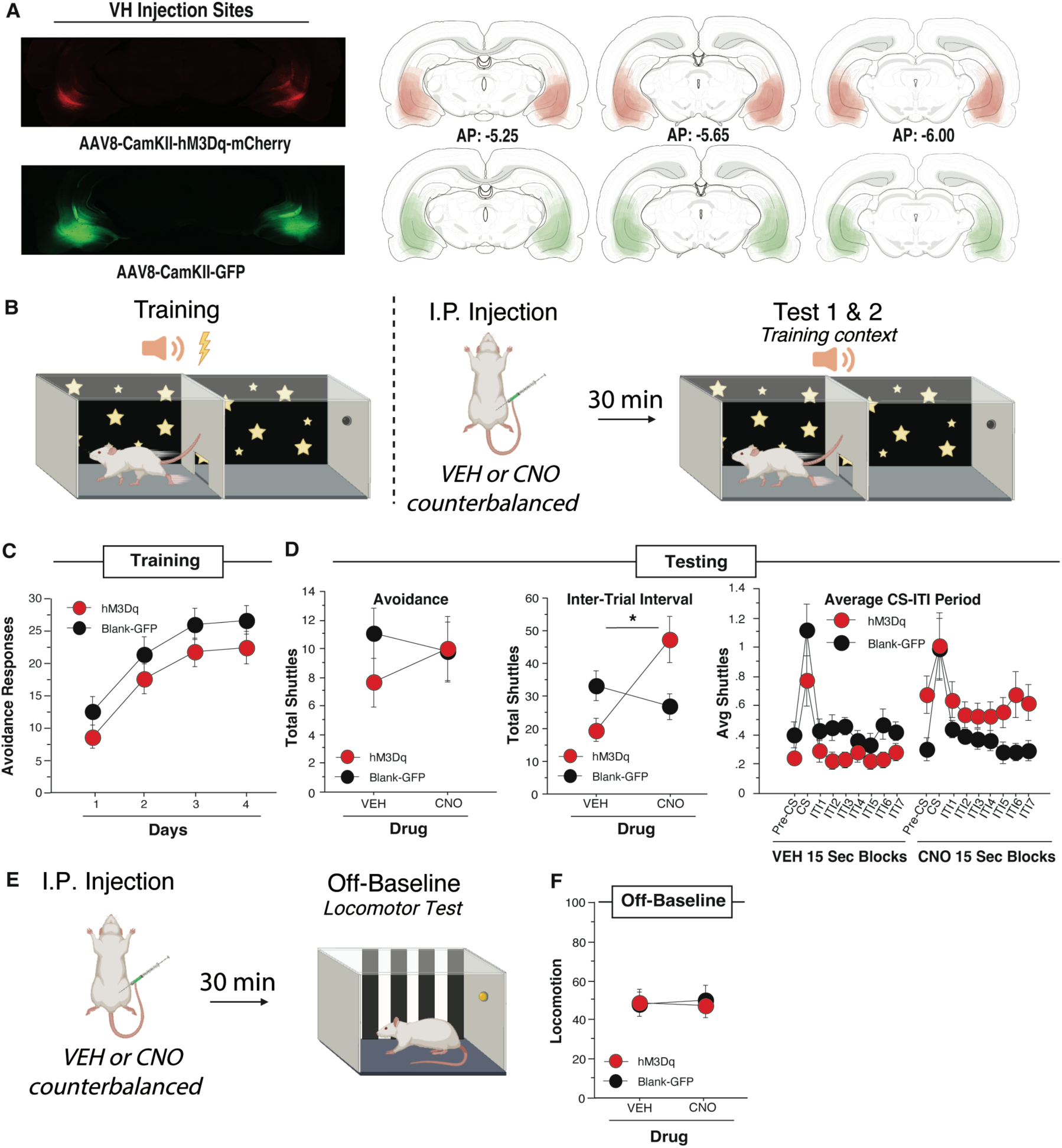
VH DREADD activation increases ITI shuttles in the training context without affecting locomotion.(A) Example virus expression and schematic viral expression (viral expression that is more common shows as darkest area) targeting vCA1 for the excitatory DREADD virus (n=14) and the blank virus (n=10). Atlas images are from Swanson, 1998 [63]. (B) Rats received 4 days of SAA training prior to counterbalanced CNO and VEH tests conducted under extinction conditions in the training context. Drug was administered with an IP injection 30 minutes prior to each test. (C) Average number of avoidance responses per day of training for active hM3Dq DREADD virus animals and blank virus-GFP animals. (D) Rats received either CNO or vehicle in an I.P. injection 30 minutes before each test; (Left) The effect of CNO and vehicle on total avoidance responses during counterbalanced drug tests; (Middle) the effect of CNO and vehicle on the total ITI shuttles over the session; (Right) Each ITI broken down into 15 second bins for a total of 8 segments for the full 2 minutes with all 10 ITI 15 second bins averaged for each animal. hM3Dq animals show a significant increase in ITI shuttles from the Vehicle to the CNO condition while blank-GFP animals do not. (E) Drug was administered I.P. 30 minutes before each locomotion test in the alternate single-chamber context. (F) Locomotion measured over a 10-minute period to assess CNO effects on movement. VH DREADD activation increases ITI shuttling without affecting acclimation, CS shuttles, or general locomotion. (* indicates p<0.05). All data presented as means ± SEM. A, B, & E created with BioRender.com

After recovery, subjects received four daily sessions of SAA training followed by two SAA tests conducted in the training context, each preceded by i.p. CNO or vehicle administered in a counterbalanced order (FIG 2B). Following SAA training, 8 subjects were determined to be poor avoiders and were excluded from the analysis (data not shown). ARs from SAA training were analyzed using a mixed-design, two-way ANOVA with a within-subjects factor of Day and a between-subjects factor of Virus (hM3Dq or Blank-GFP). This analysis revealed a main effect of Day [*F*(3, 66)=30.458, *p*<0.0001], but no main effect of virus [*F*(1, 22)=2.278, *p*=0.146] indicating that both the hM3Dq and GFP animals acquired avoidance normally (FIG 2C).

AR and ITR totals derived from SAA tests were each analyzed with a two-way, mixed-design ANOVA with a within-subjects factor of Drug (CNO or Vehicle) and a between-subjects factor of Virus (hM3Dq or Blank-GFP). There was no significant effect of VH activation on the expression of ARs at test [*F*(1, 22)=2.309, *p*=0.143] (FIG 2D left). In contrast, analysis of ITR data revealed a significant main effect of Drug [*F*(1,22)=6.06, *p*=0.0221] and a significant Virus X Drug interaction [*F*(1,22)=15.251, *p*=0.0008]. Fisher’s LSD *post-hoc* test confirmed that hM3Dq animals shuttled significantly more during the ITI in the CNO condition than in the Vehicle condition [*t*(13)=15.179, *p*=0.0016], whereas blank-GFP animals did not significantly change [*t*(9)=6.711, *p*=0.0627], demonstrating that CNO activation of hM3Dq-expressing neurons stimulated ITR expression (FIG 2D, middle).

To conduct a more nuanced analysis of VH activation during the SAA tests, each trial was divided into 15-sec time bins, beginning with a pre-CS period, continuing through the CS and into the ITI. The number of shuttles in each bin was averaged across bins within each test to generate a session-wide representation of shuttling throughout an average trial. These data were subjected to a mixed-design, three-way ANOVA with two within-subjects factors of Drug (CNO and Vehicle) and Time Bin, and a between-subjects factor of Virus (hM3Dq and Blank-GFP). This analysis revealed a main effect of Drug [*F(*1,22=5.364, *p*=0.0303] and Time Bin [*F(*8,176)=12.442, *p*<0.0001] and a Drug X Virus interaction [*F*(1,22)=14.914, *p*=0.0008]. Fisher’s LSD *post hoc* tests confirmed that hM3Dq animals showed a significant increase in shuttles when administered CNO compared to vehicle [*t*(13)=0.178, *p*=0.0016], whereas blank-GFP animals demonstrated equivalent levels of shuttling in response to both CNO and vehicle [*t*(9)=0.110, *p*=0.125] (FIG 2D, left). Thus, CNO activation of VH neurons caused a consistent increase in shuttling despite having no effect on AR totals.

Session-wide increases in the expression of shuttling could be explained by a general increase in locomotor activity caused by VH activation. To control for this possibility, a subset of subjects from this experiment (7 hM3Dq and 6 blank-GFP rats) were given a pair of locomotor tests conducted off-baseline in standard, cubicle conditioning chambers and preceded by CNO or vehicle in a counterbalanced order (FIG 2E). These data were analyzed by means of a mixed-design, two-way ANOVA with a within-subjects factor of Drug (CNO or Vehicle) and a between-subjects factor of Virus (hM3Dq or Blank-GFP), which revealed no significant effect of drug or virus [*F*(1,11)=0.830, *p*=0.382] (FIG 2F). We conclude that enhanced ITR expression caused by CNO activation of VH neurons cannot be attributed to broad increases in locomotion.

### 3.3 Experiment 3: Chemogenetic activation of VH principal neurons drives expression of ITRs in the alternate shuttle-box context

To test whether the effect of VH activation on ITRs was specific to the SAA training environment, we again expressed the excitatory hM3Dq DREADD (n=39) or GFP (n= 35) in VH principal neurons (FIG 3A). Following recovery and SAA training, all subjects were administered CNO or vehicle prior to two SAA tests conducted in an alternate shuttle-box context (FIG 3B). The poor avoider criterion was applied between training and test, and a total of 30 animals were excluded (data not shown).

**Fig. 3.**
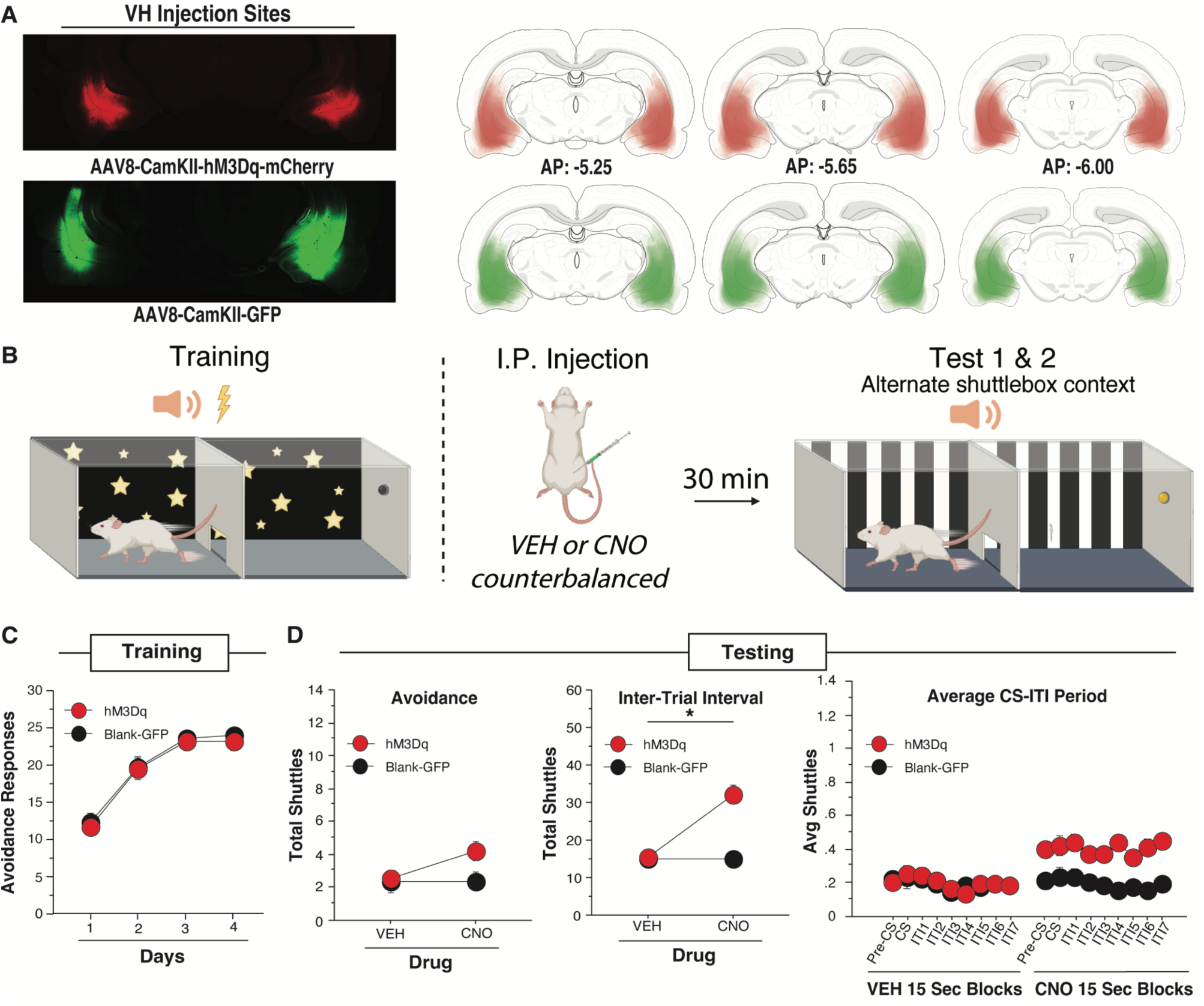
VH DREADD activation increases ITI shuttles and trends toward increasing CS shuttles in the alternate shuttle-box context.(A) Example virus expression and schematic viral expression (viral expression that is more common shows as darkest area) targeting vCA1 for the excitatory DREADD virus (n=39) and the blank virus (n=35). Atlas images are from Swanson, 1998 [63]. (B) Rats received 4 days of SAA training. Drug was administered with an IP injection 30 minutes prior to each test in the alternate shuttle-box context. A & B created on BioRender.com. (C) Average number of avoidance responses per day of training for active hM3Dq DREADD virus animals and blank virus-GFP animals. (D) Rats were then tested in various measures in the alternate shuttle-box context (Left) The effect of CNO and vehicle on total avoidance responses during counterbalanced drug tests; (Middle) the effect of CNO and vehicle on the total ITI shuttles during tests; (Right) Each ITI broken down into 15 second bins for a total of 8 segments for the full 2 minutes with all 10 ITI 15 second bins averaged for each animal. hM3Dq animals show a significant increase in ITI shuttles from the Vehicle to the CNO condition while blank-GFP animals do not. Overall hM3Dq CNO significantly increases ITI shuttles and trends toward increasing CS shuttles. (* indicates p<0.05). All data presented as means ± SEM.

Both hM3Dq and blank-GFP animals acquired avoidance normally during training. Training ARs were analyzed with a two-way repeated measures ANOVA with a within-subjects factor of Day and a between subjects factor of Virus (hM3Dq or Blank GFP) which showed no significant effect of Virus [*F*(1,72)=0.126, *p*=0.724]. This analysis did reveal a main effect of Day [*F*(3, 216)=81.286, *p*<0.0001], indicating that all animals improved over time (FIG 3C).

Total ARs performed during the SAA tests in the alternate shuttle-box context were analyzed using a mixed-design, two-way ANOVA with a within-subjects factor of Drug (CNO or Vehicle) and a between-subjects factor of Virus (hM3Dq or Blank-GFP). This revealed a main effect of Drug on ARs [*F*(1,72) = 4.757, *p* = 0.0325] and nonsignificant trend toward a Virus by Drug interaction [*F*(1,72) = 3.872, *p* = 0.053), indicating that CNO activation of VH neurons produced a marginal increase in AR expression (FIG 3D, left). However, an identical ANOVA on ITR data derived from the same tests revealed a very significant Virus X Drug interaction [*F*(1,72) = 32.178, *p* < 0.0001], as well as a main effect of Virus [*F*(1,72) = 11.128, *p* = 0.0013] and a main effect of Drug [*F*(1,72)=31.744, *p* < 0.0001]. This was confirmed with Fisher’s LSD *post-hoc* test that showed that hM3Dq animals shuttled significantly more during the ITI in the CNO condition than in the VEH condition [*t*(38)=4.905, *p*<0.0001] while control-GFP animals did not significantly change [*t*(34)=3.276, *p*=0.972]. When sex was added as a factor to each of these ANOVAs, there was a main effect of sex on ARs [*F*(1,70) = 5.066, *p* = 0.0275] and ITI shuttles [*F*(1,70) = 5.295, *p* = 0.0244] demonstrating that female rats generally shuttle more in the alternate shuttle-box context than male rats regardless of virus or drug. In summary, VH activation significantly increased ITR expression and trended toward increasing ARs in the alternate shuttle-box context (FIG 3D, middle).

To perform a more nuanced analysis of shuttling across each test session, we again broke each trial into 15-sec epochs and generated session-wide averages of shuttling within each epoch, as described under Experiment 2. These data were analyzed with a mixed-design, three-factor ANOVA with two within-subjects factors of Drug (CNO or Vehicle) and Time Bin and a between-subjects factor of Virus (hM3Dq or Blank-GFP). This analysis revealed a main effect of Virus [*F(*1,72)=9.230, *p*=0.0033], Drug [*F(*1,72)=29.907, *p*<0.0001], and Time Bin [*F(*8,576)=2.788, *p*=0.0049], as well as a Virus X Drug interaction [*F*(1,72)=29.539, *p*<0.0001]. Fisher’s LSD *post hoc* tests confirmed that hM3Dq animals showed a significant overall increase when given CNO compared to vehicle [*t*(38)=0.061, *p*<0.0001], whereas Blank-GFP animals showed equivalent responding in response to both CNO and vehicle [*t*(34)=0.044, *p*=0.977] (FIG 3D, right). Thus, CNO activation of VH neurons produced a uniform increase in shuttling that remained flat across the session, instead of selectively increasing shuttles at any point, including during the CS. These data are consistent with the interpretation that VH activation is sufficient to drive ITR expression in both training and alternate shuttle-box contexts.

### 3.4 Experiment 4: Context Extinction does not significantly reduce ITI shuttling

Because we hypothesized that an aversive association between context and the US is a key associative structure underlying ITRs, we examined the effect of contextual extinction via prolonged exposure on ITR expression. All subjects received SAA training before being broken into one of four conditions that received either: exposure to the SAA context (n=13), exposure to an alternate chamber and 30 CSs/day (n=14), exposure to an alternate chamber and no CSs (n=14), or exposure to a five-gallon bucket (i.e. handling control, n=14). These manipulations were followed by an SAA test in the training context for all subjects (FIG 4A). The poor avoider criterion was applied between the training and extinction phases, and 5 animals were excluded. Training ARs were analyzed using a two-way repeated-measures ANOVA with a within-subjects factor of Day and the between-subjects factor of Exposure Group. This ANOVA revealed a main effect of Day [*F*(3,153) = 160.007, *p* <0.0001], but no effect for Exposure Group [*F*(3,51) = 0.585, *p* = 0.628] indicating that all groups acquired avoidance normally (FIG 4B, left).

**Fig. 4.**
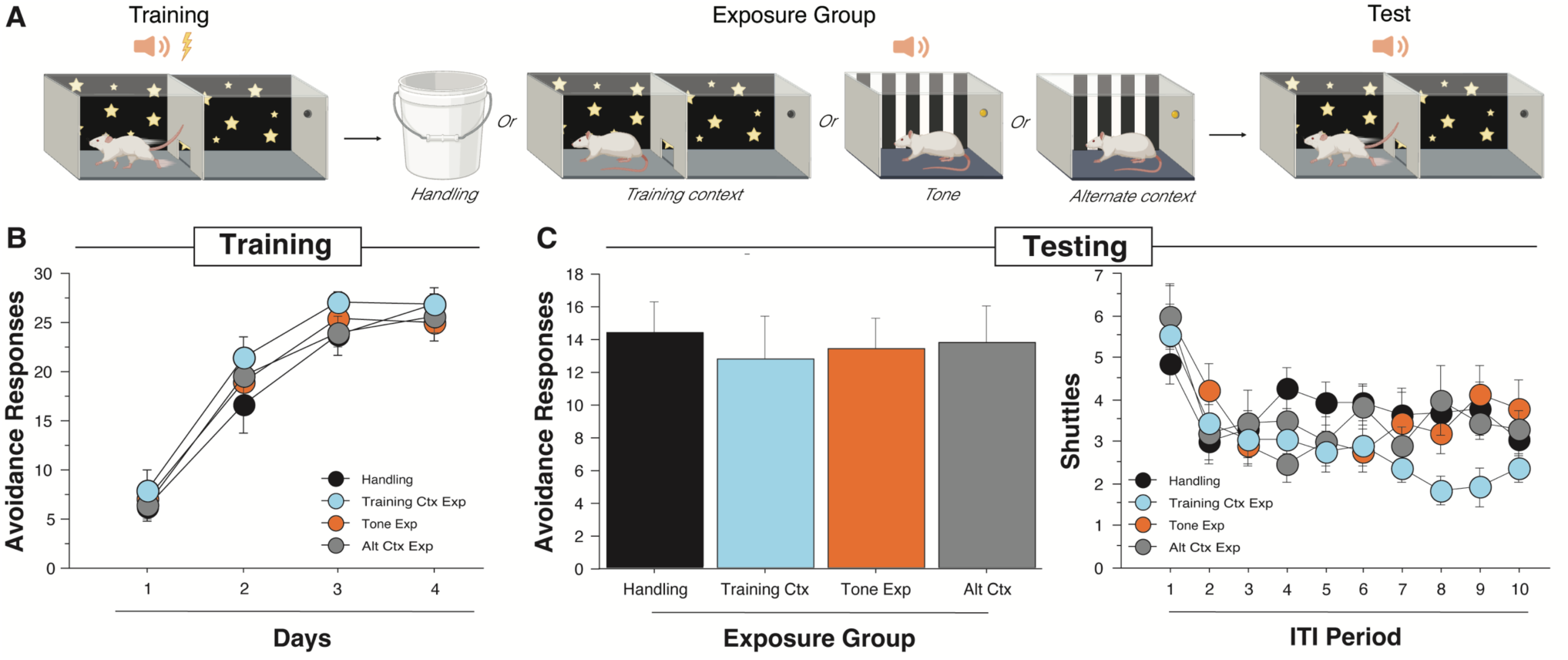
Context extinction does not significantly reduce ITRs.(A) Rats received 4 days of two-way SAA training. Rats were then assigned to groups where they were either exposed to the training context, the alternate chamber, the alternate chamber with the avoidance tones presented, or equivalent handling for the same period as typical avoidance training for 3 days. They finally received a 10-tone test under extinction conditions on day 8 to assess their avoidance and ITI shuttling behavior. Created with Biorender.com (B) Average avoidance responses performed each day of training. Rats were assigned to groups so that each groups would have similar levels of avoidance in the handling (n=14), training context (n= 13), tone exposure (n=14), and alternate context (n=14) groups across the days. (C) (Left) Total avoidance responses in the test for each group. All exposure groups showed similar levels of CS shuttling. (Right) Number of shuttles for each ITI throughout the 10-tone test. Training context extinction marginally but not significantly decreases ITI shuttling during the test compared to the other 3 control groups. All data are shown as mean ± SEM.

AR totals from the SAA test were analyzed using a one-way ANOVA that compared the four exposure groups. This revealed no effect for group [*F*(3,51) = 0.096, *p* = 0.962], demonstrating that none of the exposure manipulations influenced the robust expression of avoidance (FIG 4C, left). We also analyzed ITR expression across the 10 ITIs of the test session using a two-way, repeated-measures ANOVA with a within-subjects factor of ITI Period and a between-subjects factor of Exposure Group. This revealed a main effect of ITI period [*F*(9,459)=9.853, *p*<0.0001], indicating that ITRs decreased over time in all groups. However, the ITI X Exposure Group interaction was nonsignificant [*F*(27, 459) = 1.431, *p* = 0.0763] (FIG 4C, right), demonstrating that ITR behavior did not differ between animals that underwent context extinction and our control manipulations (FIG 4C, right). Further, we note that the overall pattern and level of ITR expression across the SAA test session is roughly typical of what we observe in subjects that have not received any additional manipulations between training and test (see: FIG 1B, right for comparison). We conclude that context fear may contribute to ITR performance but is not the primary driver of this action.

## 4 Discussion

Here, we report that pharmacological inhibition of ventral hippocampus (VH) reduces the expression of inter-trial responses (ITRs – shuttling during the inter-trial interval) in a two-way signaled active avoidance (SAA) paradigm. In addition, we demonstrate that chemogenetic excitation of principal neurons in VH is sufficient to enhance the expression of ITRs when tested in both training and alternate shuttle-box contexts. Neither manipulation had an effect on avoidance responses (ARs – shuttling during the CS), and control experiments revealed no effect of VH activation on locomotion, suggesting that VH selectively facilitates ITRs when activated, instead of producing an overall increase in locomotor activity. Though we hypothesized that context-shock associations would be a critical associative substrate for ITR expression, extinction of the training context via prolonged exposure had no effect on ITRs. We conclude that context-shock associations may still play some role, but that VH is primarily engaged in some other associative/motivational process to drive the expression of ITRs.

The hypothesized role for context-shock associations in VH-mediated ITRs was grounded in work demonstrating that VH neurons display place fields [29–32] and respond to presentation of a foot-shock US [11,33]. Indeed, VH has been shown to mediate multiple aversive/associative contextual processes [8–15,17,18]. Particularly relevant to this study is the demonstration that contextual extinction via prolonged exposure to the training environment reduces ITRs in a wheel-run signaled avoidance task [6]. For all of these reasons, we were surprised to find that extinction of context-shock associations had no effect on VH-mediated ITRs in the two-way SAA paradigm. However, it has been demonstrated that context exposure reduces ITRs in Roman high-avoidance rats bred for augmented performance in SAA (i.e. enhanced expression of both ARs and ITRs), suggesting that an aversive association between the US and the training context may indeed account for the degree to which Roman high-avoidance rats differ from Sprague-Dawley rats (the strain employed here) in ITR expression [34]. Thus, context-shock associations could function to boost ITRs in subjects with a genetic propensity for high avoidance, even if they are not comparably operative in subjects lacking the same inborn tendencies. Further, though superficially similar, behavior in distinct avoidance tasks (i.e. wheel-run vs two-way SAA) may rely on distinct forms of aversive memory.

Associative structures other than direct associations between the training context and shock may play a role in the generation of ITRs in two-way SAA. Contexts can serve as occasion setters and modulate both Pavlovian and instrumental behavior without direct associations with USs, CSs, or the responses they evoke [35]. Several studies have shown that the hippocampus has an important role in occasion setting [36,37], including in procedures in which contexts serve as occasion setters [38,39]. Although most studies have focused on the role of the dorsal hippocampus in occasion setting, VH does play a role in this process, including the context-dependent modulation of Pavlovian extinction [13]. Hence, it is possible that the VH is recruited to produce ITRs via the occasion-setting properties of the shuttle-box context, rather than direct context-shock associations.

Though manipulations of context often involve the physical apparatus in which behavior is acquired and tested, a comprehensive consideration of contextual process should include any aspect of the background against which action occurs, including psychological processes intrinsic to the subject [37]. It has been argued that a residual aversive state triggered by CS presentation is an element of the interoceptive context that energizes ITR expression [25]. Indeed, SAA extinction via presentation of the CS in the absence of the US leads to the suppression of both ARs and ITRs [40], demonstrating that CS-evoked aversion may contribute to both responses. Our results may thus suggest that a lingering cue-elicited aversive state can engage VH, which then transforms this interior state into the behavioral output of ITRs via the recruitment of the appropriate efferent pathway, consistent with recent conceptual frameworks describing VH function [24]. From a preclinical perspective, these VH-mediated responses can be considered perseverative behaviors decoupled from any exterior reinforcement that provide a viable model of the behavioral compulsions observed in obsessive compulsive disorder (OCD; [41,42]). Indeed, recent brain imaging work in human patient populations links hippocampal volume and functional connectivity to OCD symptomology [43,44]. VH-mediated ITRs may thus serve as a valuable research tool with which to generate mechanistic insight into compulsive behavior and thus multiple, related psychopathologies.

Other VH-mediated processes may explain our results. All SAA tests reported here were conducted under extinction conditions, meaning that, unlike training, the CS was presented for a full 15-seconds regardless of whether the subject produced an AR. Previous work indicates that VH maintains behavioral responses when CSs are presented for longer than expected [45] and mediates behavioral responses under other forms of timing uncertainty [46,47]. Furthermore, it has been shown that VH activation can increase context generalization [48], which may explain the observation that VH activation drives up ITR expression in both the training and alternate shuttle-box contexts. Finally, VH lesions have been linked to perseverative responding in other tasks[49,50]. Because strong forms of neuronal activation can dysregulate hippocampal function [48,51], it is possible that DREADD excitation of VH principal neurons produced a deficit in VH function that released perseverative responding to the CS in the form of ITRs.

The data presented here, in conjunction with a previous report demonstrating that VH suppresses ARs in an alternate shuttle-box context [18], suggest that VH supports dissociable behavioral functions in SAA. VH sends projections to multiple avoidance-relevant brain areas, including the basolateral amygdala [15,26,52–54], nucleus accumbens [16,52,54–56], infralimbic cortex [14,56,57], and the bed nucleus of the stria terminalis [52,58]. Inputs from VH can either stimulate or suppress the activity of principal neurons within the same target region, via direct excitatory connections or feedforward inhibition, respectively [59–62]. This complex circuit architecture may explain the nuanced effect of VH manipulations on shuttling behavior in SAA, facilitating these responses in the case of ITRs and exerting a context-specific form of suppression in the case of ARs [18]. Future work will explore the hypothesis that these contrasting effects are supported by the differential recruitment of distinct efferent projections arising in VH.

## CRediT authorship contribution statement

**Cecily R. Oleksiak:** Investigation, Methodology, Formal analysis, Validation, Visualization, Writing-original draft. **Samantha L. Plas:** Investigation, Formal analysis. **Denise Carriaga:** Investigation. **Krithika Vasudevan:** Investigation. **Stephen Maren:** Conceptualization, Methodology, Resources, Visualization, Supervision, Project administration, Funding acquisition, Writing-review & editing. **Justin M. Moscarello:** Conceptualization, Methodology, Resources, Visualization, Supervision, Project administration, Funding acquisition, Writing-review & editing.

## Declaration of interest

none

## Funding

This work was supported by the National Institute of Mental Health [RO1MH065961]

## Notes

### Competing Interest Statement

The authors have declared no competing interest.

